# Native extracellular vesicles display surface bound RNAs that are co-delivered to cells

**DOI:** 10.64898/2025.12.17.694909

**Authors:** Hannah Weissinger, Madhusudhan Reddy Bobbili, Yan Yan, Maria Gockert, Elsa Arcalis, Johannes Grillari, Mette Galsgaard Malle, Jørgen Kjems

**Author notes:** Corresponding authors, for M.G.M and for J.K.

## Abstract

Extracellular vesicles (EVs) can transport functional RNA between cells and therefore hold great potential for diagnostics and RNA-based therapeutics. Classically, RNA is believed to be encapsulated in the EV lumen. However, it has recently been demonstrated that cells present RNA on their surface. This RNA was found to be glycosylated, and although glycosylated tRNA was also found in EVs, its exact location remained elusive. Here, we demonstrate the presence of RNA on the surface of mesenchymal stem cell (MSC) derived EVs. By combining single-vesicle measurements with direct and selective visualization of RNA, we introduce surface RNA (surfRNA) as a new inherent component of EVs. RNA sequencing supports the surface localization of this RNA and further identifies tRNA fragments as primary constituent of surfRNA. Importantly, surfRNA is co-delivered to target cells together with EVs, suggesting a yet unrecognized uptake route of extracellular RNA. A deeper understanding of the surface-associated RNA may have significant implications for EV biogenesis, targeting, and downstream functional effects. We further envision that these findings are transferable to other nanoparticles and will thereby advance the field of therapeutic RNA delivery.

## Introduction

RNA is fundamental in directing cellular function. Intracellular, it performs a multitude of functions, among them the most crucial processes of life. The role of extracellular RNA (exRNA) is less well understood, but RNA is known to be transported through the extracellular space and taken up into cells. This phenomenon occurs naturally in the case of cellular release and internalization of viruses^1, 2^ or extracellular vesicles (EV)^3–6^ internalization, but is also utilized strategically to deliver therapeutic RNAs using synthetic lipid nanoparticles (LNPs)^7, 8^ or EVs^9–11^. EVs are cell-derived nanoparticles that within a surrounding lipid membrane contain a large variety of biomolecules including miRNAs, short mRNAs and mRNA fragments, tRNAs and tRNA fragments (tRFs), as well as DNA, protein, and metabolites^12^. EVs are present in all biofluids and can deliver their content to target cells through fusion with the cell membrane or uptake into the cell. Similar to viruses and LNPs, EVs are surrounded by a biomolecular corona^13–15^. The corona consists of proteins that, upon exposure to biofluids, adhere to the particle surface. For EVs, the corona has been shown to impact biodistribution, immune detection, and stability^16–18^.

RNA encapsulation into EVs, LNPs or other membrane particles offers RNA protection from enzymatic degradation by RNases, stabilizing it in the extracellular environment. Alternatively, RNA can gain stability by forming structures, one example being tRFs that resist RNases by forming dimers^19, 20^. Recent findings further suggest the presence of RNA on the surface of cells and propose glycosylation as a mechanism to protect it^21^. GlycoRNA has recently been found associated with EVs by two studies^22, 23^. However, these findings are contradictory concerning the exact localization of glycoRNA, leaving unclear whether RNA is present on the EV surface or solely inside the lumen. EVs are a heterogeneous mixture of vesicles, as even single cells can release different types of vesicles. In body fluids, an additional complexity is added by the presence of EVs of various cell types. A more thorough understanding of EVs and their RNA can only be achieved by single-vesicle analysis to deconvolute the heterogeneity in EV size, surface RNA density and to directly distinguish surface RNA from potential co-isolated RNA in the EV solution. Further, advancing the knowledge about the identity of exRNA and EV-associated RNA is crucial for their future use as a diagnostic tool^24^.

Here, we use a variety of single-vesicle measurements to directly show the presence of RNA on the surface of EVs, which we term surfRNA (surface RNA). We isolate and analyze EVs derived from human Wharton’s Jelly mesenchymal stem cells (WJ-MSCs) immortalized by telomerase, resulting in a continuously growing primary like cell line, with high therapeutic relevance^25^. At the single-vesicle level, we confirm the presence of surfRNA using RNA specific fluorescent labelling and visualization techniques combined with nanoparticle flow cytometry and total internal reflection fluorescence (TIRF) microscopy. In addition, in a complementary small RNA sequencing approach, surfRNAs are identified to mainly be tRFs. We further observe, using live-cell imaging, that surfRNA colocalizes with EVs when entering cells, suggesting an undiscovered cellular import pathway for exRNA. The novel insights into the interaction of EVs and RNA on the nanoscale may be relevant to research on RNA on the surface of cells. Understanding the nucleic acid association with EVs will be a crucial step towards their application in drug development and diagnostics.

## Results

### SurfRNA detection by fluorescent poly(A)-tailing and nanoparticle flow cytometry

To investigate the localization of EV-associated RNA, we studied WJ-MSC-derived EVs that were isolated by tangential flow filtration (TFF), a gentle and scalable method based on the particle size^26^. EVs were characterized by nanoparticle tracking analysis (NTA), western blot, transmission electron microscopy (TEM), and bead-based multiplex analysis according to the MISEV guidelines^27^ and exhibited size, shape and surface markers characteristic for EVs (Supplementary Fig. S1). To visualize the RNA, we developed a sensitive labeling method suitable for readout by nanoparticle flow cytometry. Detection of RNA in single EVs often relies on knowledge about the sequence and the ability of a detection probe to basepair with the target RNA^28, 29^. To detect EV-associated RNA of unknown identity, detection methods must rely on shared RNA properties. We therefore established an assay based on poly(A) tailing with fluorescently labeled adenosine triphosphate (ATP). The employed enzyme Poly(A) polymerase, E coli (E-PAP) has a high specificity for RNA substrates and polyadenylates 3’ termini of RNAs with multiple fluorescent adenosine residues^30^. The resulting amplification of fluorescent signal enhances the sensitivity to detect even low-abundant RNA on the outside of the EV whereas the interior RNA is separated from the enzyme by the EV membrane (Fig. 1a).

**Fig. 1.**
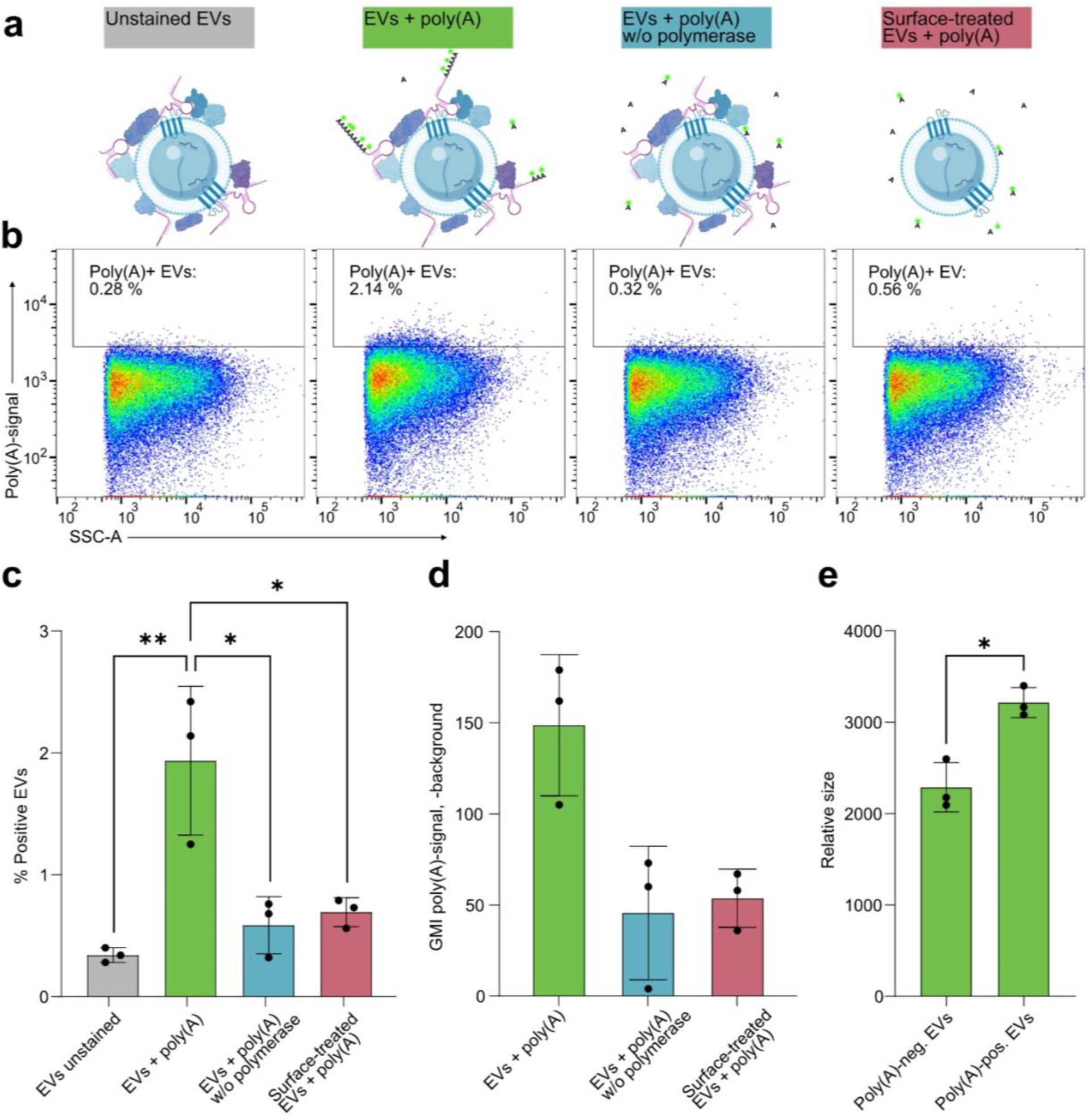
Accessibility for polyadenylation shows the localization of RNA on the surface of Extracellular Vesicles (EVs). Nanoparticle flow cytometry was used for single-particle detection of EVs. **a** Schematic of EVs with polyadenylated surface RNA and experimental controls. **b** Scatterplots showing the percentage of polyadenylated EVs in one representative experiment. Side scatter area (SSC-A) signal vs poly(A) signal. **c** Percentage of poly(A)-positive EVs as gated in B for three independent experiments. Individual datapoints with mean and standard deviation are depicted. RM one-way ANOVA with Tukey’s multiple comparisons test. **d** Quantification of the geometric mean intensity of the EVs for the poly(A) signal in three independent experiments. Individual datapoints with mean and standard deviation are depicted. **e** Comparison of relative EV size (geometric mean of SSC-A) of positive and negative populations for polyadenylation. Individual datapoints with mean and standard deviation are depicted for three independent experiments and compared with Wilcoxon test.

The fluorescence of individual EVs was then sensitively detected using nanoparticle flow cytometry. A subset of about 2 % of the EVs were found to be fluorescently labeled by polyadenylation, indicating the presence of RNA and available 3’ ends on the surface (Fig. 1b and 1c). To exclude that the signal stems from nonspecific binding of fluorescently labeled ATP to EVs, we incubated EVs with the labeled ATP nucleotides without adding polymerase (EVs + Poly(A) w/o polymerase). As a second control, we surface treated EVs enzymatically with Proteinase K and RNase A to remove the EV corona and surfRNA (Surface-treated EVs + poly(A)). In both controls, the percentage of labeled EVs was significantly reduced compared to EVs with poly(A) treatment (EVs + Poly(A)).

In addition to fluorescence, nanoparticle flow cytometry simultaneously detects particle size at the single EV level from violet side scatter (vSSC) detectors. In the polyadenylated samples, the polyadenylated EVs were approximately 40 % larger compared to EVs that did not show polyadenylation-induced fluorescence, probably contributed by the protruding RNA brush created by the enzyme (Fig.1d). Detergent-mediated lysis of EVs led to a decrease in particle number in the EV gate and to an increase in small particles as expected (Supplementary Fig. S2b).

Co-staining EVs with a fluorescent membrane stain showed similar polyadenylation-derived fluorescence increase in the membrane-stained EV population, confirming that the detected signal stems from membrane particles, not protein aggregates of similar size (Supplementary Fig. S2c). Performing the poly(A) reaction and membrane staining in PBS without EVs only caused a neglectable background signal (Supplementary Fig. S2c).

### Direct labeling reveals nucleic acids on a larger subset of EV

Labeling RNA by polyadenylation requires an accessible 3’-hydroxyl group^31^. Considering that 3’ ends of surface-associated RNA could be hidden by RNA structure, proteins or chemical modifications, we applied a complementary labeling approach independent of the RNA ends. TOTO-3 is a dimeric cyanine nucleic acid stain that is membrane impermeable and undergoes significant fluorescence enhancement upon binding to nucleic acids. To test its suitability for detecting surfRNA, we again used WJ-MSC-derived EVs. To analyze TOTO-3-stained EVs, total internal reflectance fluorescence (TIRF) microscopy was chosen due to its superior sensitivity. EVs were stained with TOTO-3 and a lipidated membrane dye, immobilized on passivated glass slides and imaged in a semi-automatic fashion to capture hundreds of fields of view (FOV) without human-biased selection (Fig. 2a). Individual EVs were detected, and their fluorescence signals were integrated in the acquired FOVs using an in-house developed image analysis algorithm. The colocalized membrane signal with the intensity of the TOTO-3 signal was quantified for each individual EVs (Fig. 2b and Supplementary Fig. S3b). Using an automated signal threshold of 2.5 x standard deviation of the TOTO-3 RNA signal in the negative control of each experiment, 9.1% ± 0.06 of the EVs were detected positive for TOTO-3 staining (Fig. 2c and d). Surface treatment with Proteinase K and RNase A caused a significant reduction in the percentage of positively stained EVs in agreement with surface localization of the RNA. The relative particle size can be obtained from TIRF microscopy data through its correlation with the membrane labeling intensity^32, 33^. Comparing TOTO-3-positive and TOTO-3-negative EV populations indicated that the presence of RNA on the EV surface was independent of the particle size (Fig. 2e). Testing for correlation between membrane signal and TOTO-3 signal, the Pearson’s correlation coefficient was <0.06 in all three repetitions, confirming the lack of correlation between EV size and surfRNA amount. To verify that the TOTO-3 dye is membrane-impermeable and cannot stain RNA inside the EV lumen, we used liposomes as a model system. Liposomes that contained tRNA in the lumen and empty liposomes were stained with TOTO-3 and measured by TIRF microscopy. While a clear fluorescence shift was visible for EVs, no difference in fluorescence was found for liposomes, confirming the strict exclusion of the dye from the liposomal interior (Fig. 2f and 2g and Supplementary Fig. S4).

**Fig. 2:**
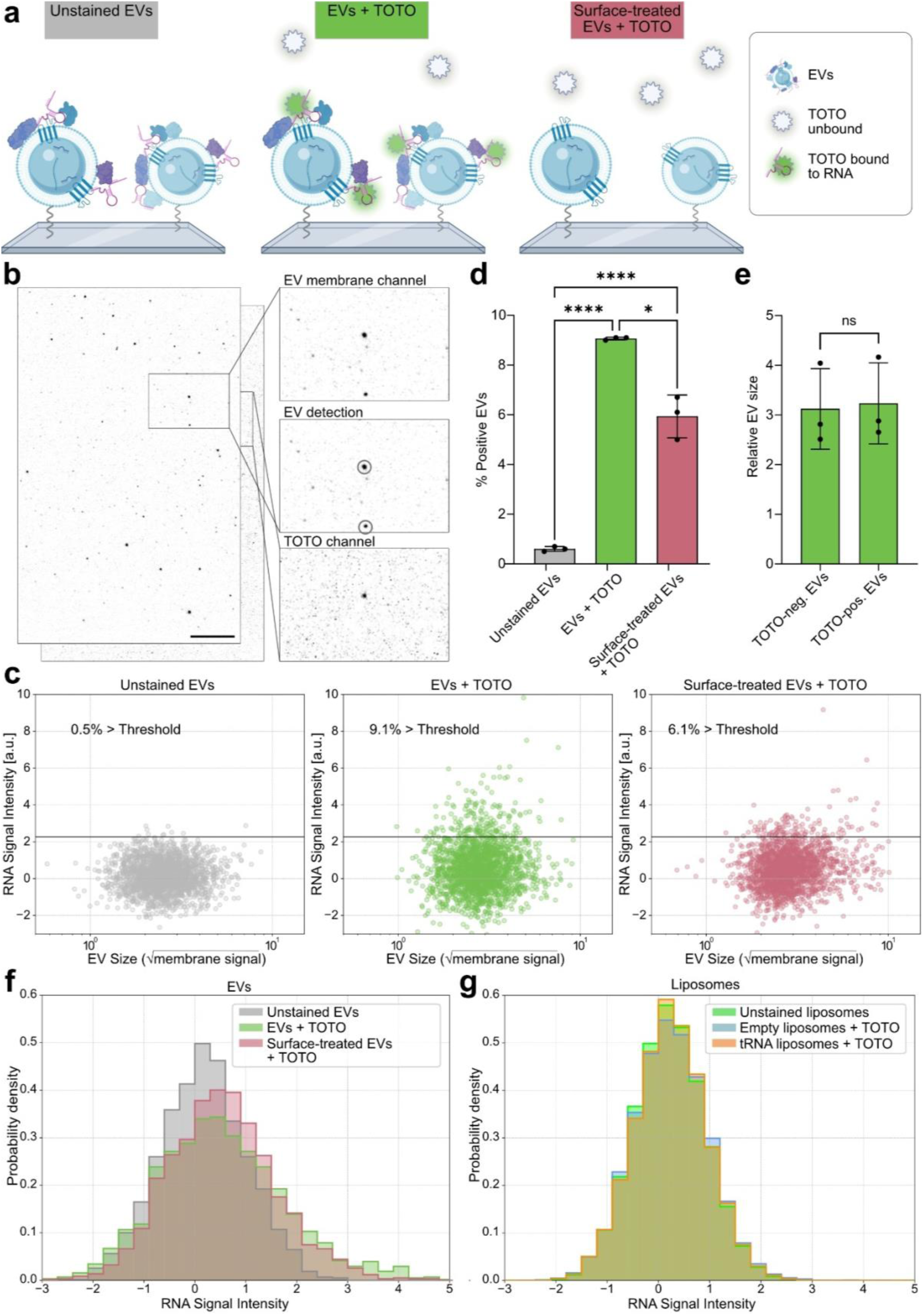
Direct staining confirms the presence of external nucleic acids. **a** Schematic of surfRNA detection by TIRF microscopy. **b** Representative field of view showing the co-localization of membrane signal and TOTO-3 signal. EVs were detected by automated image analysis and the signal within the black circle was integrated for each spot. Scale bar = 10 µm. **c** Scatterplots showing the percentage of TOTO-3-stained EVs in one representative experiment. Number of analyzed EVs: n=1960. **d** Percentage of TOTO-3-positive EVs for three independent experiments. Individual datapoints with mean and standard deviation are depicted. RM one-way ANOVA with Tukey’s multiple comparisons test. **f** Histogram showing the TOTO-3 signal intensity in EVs. **g** Histogram showing the TOTO-3 signal intensity in liposomes with and without encapsulated tRNA.

### RNA sequencing reveals the identity of surfRNA

Having confirmed the presence of surfRNA through two independent single-particle analysis methods, we next sought to identify the types of RNA present on the EV surface. As EVs mostly contain short RNAs, small RNA sequencing is a well-established method to study EV-associated RNA^34, 35^. However, it is a bulk method which cannot distinguish between RNA attached to EVs and free RNA that could be co-isolated as an impurity. While TFF is a gentle isolation method that has been previously used for EV corona studies^36^, it enriches all particles of a certain size. In contrast, affinity isolation more specifically isolates EVs with a certain marker and was therefore chosen for the first sequencing approach. The recently developed Snorkel-tag is a versatile and specific affinity isolation method, based on a fusion construct of HA-tag and CD81 and bead-based isolation of HA-positive EVs in vitro and in vivo^37^. In addition to WJ-MSC-derived Snorkel isolated EVs (HA-tag EVs), we included samples isolated by direct CD81 immunoprecipitation, including unmodified/WT WJ-MSCs (CD81 WT EVs), WJ-MSCs stably expressing the Snorkel construct (CD81 S7 EVs) or with a control construct (CD81 E3 EVs).

To distinguish between extravesicular RNA and RNA encapsulated in the EV lumen, we performed RNA sequencing of the enriched WJ-MSC EVs comparing native, untreated EVs with EVs that were treated with Proteinase K and RNase A. Further, we included samples of EVs that were lysed and subsequently treated with Proteinase K and RNase A. Principal component analysis (PCA) showed that the surface treatment was the main factor to distinguish between the samples, indicating a different RNA profile of extravesicular RNA compared to RNA in the EV lumen (Fig. 3a and Supplementary Fig. S5a). We then directly compared untreated and surface-treated samples from the same batch of EVs. Interestingly, in all four complementary isolation-technique groups specific tRFs were more abundant in untreated samples (Fig. 3b and Supplementary Fig. S3c). Strikingly, tRFs that were significantly enriched on the surface were overlapping between the isolation techniques to a high degree (Fig. 3c). As expected, there was slightly more overlap between the three CD81-based isolation techniques than between those and the HA-tag isolation. In contrast, miRNAs and other small RNAs were only marginally affected by the surface-treatment (Supplementary Fig. S5d and S5e).

**Fig. 3:**
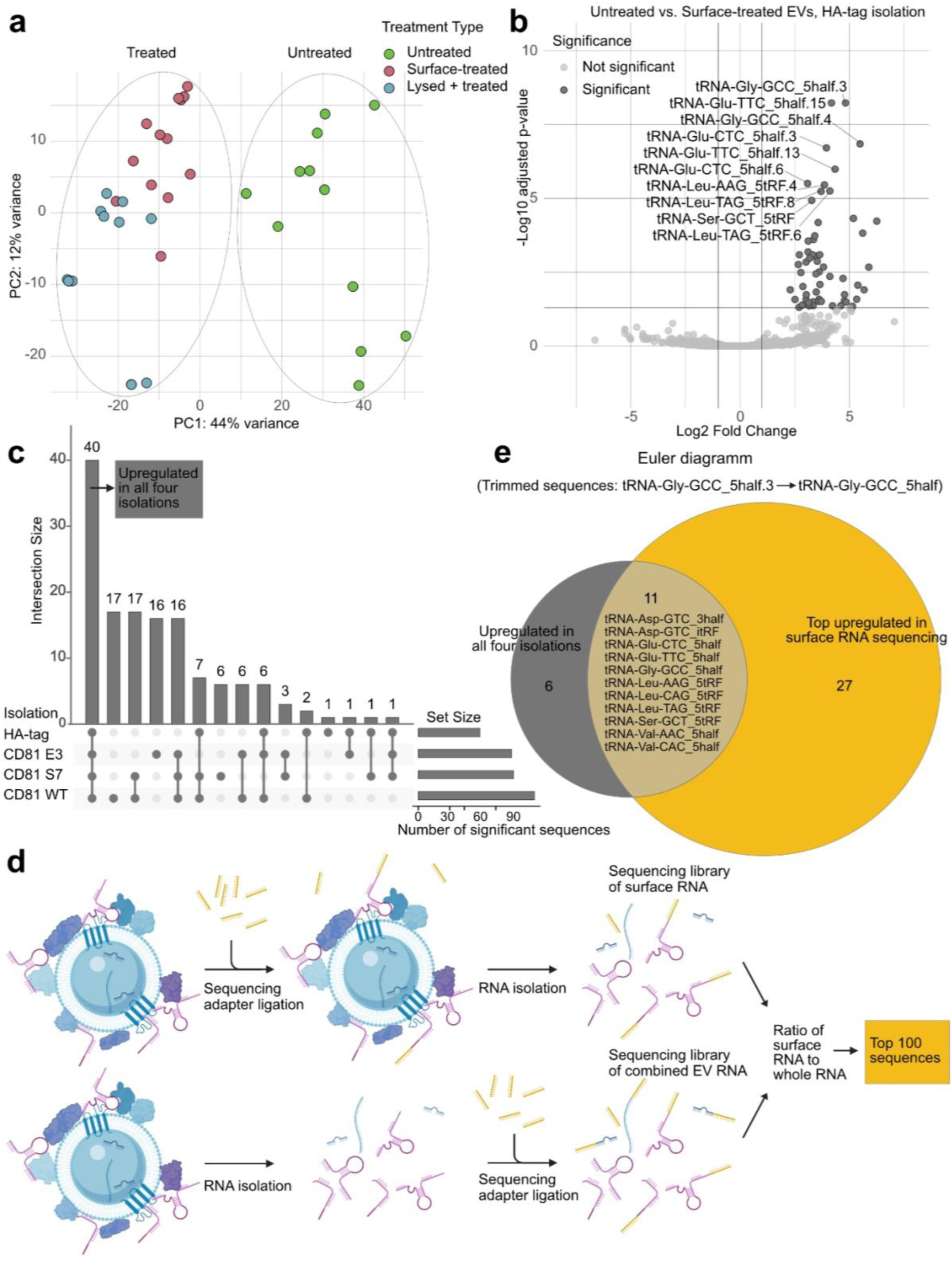
Specific tRFs are enriched on the surface of Extracellular Vesicles. **a** Principal component analysis (PCA) of variance-stabilized tRFs of untreated EVs, surface-treated EVs (Proteinase K + RNase A) and lysed and treated EVs. **b** Volcano plot for tRFs of EVs isolated by HA-tag, comparing untreated and surface-treated EVs. Significance was defined as FDR < 0.05 and |log₂ fold change| > 1. The 10 sequences with the lowest adjusted p-value are labeled. **c** UpSet plot comparing the sequences that were significantly upregulated on the surface across four isolations. **d** Strategy for selective sequencing of surface-RNA. Sequencing adapters were ligated on intact EVs to exclude encapsulated RNA. As a control, whole EV RNA was sequenced by isolating EV RNA and subsequent adapter ligation. **e** Euler diagram showing the overlap of tRFs that were surface-enriched in all four isolations indicated in panel c with the 100 most upregulated sequences of the selective surface RNA sequencing illustrated in d. Sequences were trimmed and compiled into 3’-/5’-half/internal fragment classes to allow for differences in sequence length, causing a reduction to 17 shared sequences across isolations and 38 sequences derived from the selective surface RNA sequencing.

Next, we wanted to confirm those results and connect them to the single-particle analysis. As both nanoparticle flow cytometry and TIRF microscopy experiments were performed using TFF-isolated EVs, we now aimed to obtain the sequences of their surfRNA. Based on the observation that the 3’-ends of the RNA were accessible for polyadenylation, we hypothesized that a sequencing adapter could be ligated to them without prior isolation of the EV RNA. This would result in a sequencing library exclusively consisting of surfRNA. In parallel, we isolated EV RNA prior to adapter ligation and produced a combined library of surfRNA and interior RNA (Fig. 3d). Following small RNA sequencing, we quantified the relative amount of individual surface RNA. The 100 most surface-enriched sequences (≥19 fold enrichment), were then compared to the 40 sequences that were consistently and significantly upregulated in the affinity purified, untreated EVs vs Proteinase K and RNase A treated EVs (Fig. 3c). As many tRFs are highly modified, premature termination of reverse transcription can introduce biases during library preparation^38^. To create a more robust comparison independent of the sequencing approach, we chose to compare sequences by their anticodon and identity (5’ half or 5’ fragment, 3’ half or 3’ fragment or internal fragment) instead of their exact sequence. This caused a reduction from 40 to 17 sequences for the dataset comparing untreated EVs with EVs treated with Proteinase K and RNase A and from 100 to 38 sequences for the dataset based on external adapter ligation, with an overlap of eleven sequences between the two datasets (Fig. 3e). Interestingly, we found the list to be dominated by 5-halves of 5’ fragments. The comparison of the exact sequences resulted in an overlap of 5 sequences and is shown in Supplementary Fig. S5b. Collectively, these findings suggest that surfRNA on WJ-MSC-derived EVs is predominantly composed of tRFs, the majority of which correspond to 5’-derived fragments.

### Live cell imaging shows simultaneous delivery of EVs and surfRNA to recipient cells

EVs are internalized by target cells through mechanisms such as endocytosis and membrane fusion^39–41^. The delivery of the EV cargo has been studied extensively^3, 42, 43^. However, surfRNA is located outside of the vesicle and the fate of surface-bound molecules during cellular uptake of EVs remains poorly understood. We therefore set out to investigate whether the surfRNA can be co-delivered to target cells. We again visualized surfRNA by polyadenylation with fluorescently labeled ATP. Membrane stained and polyadenylated EVs were incubated with HeLa cells for 1 h, prior to live cell imaging by spinning disc confocal microscopy. The surfRNA (green signal) predominantly appeared colocalized with the EV membrane signal (red signal) (Fig. 4a). Interestingly, EVs and surfRNA were detected both in the periphery of the cell and in the perinuclear region, indicating that surface-bound RNA can enter cells along with EVs (Fig.4a and b – see representative EV i). However, we also observed surfRNA distant from the nucleus, without corresponding EV membrane signal, which suggests EVs fusion with the cell membrane, leaving the RNA behind on the cell surface (Fig.4a – see representative EV ii). More examples are presented in additional fields of view (Supplementary Fig. S6c). In control EVs without polyadenylation, no signal was observed in the respective channel, while EV membrane signal was unaltered. Additional negative controls with membrane staining and polyadenylation in PBS with no EVs present exclude direct labeling of cellular structures (Supplementary Fig. S6a). Z-stacks further confirm the co-localization of EVs and surfRNA in close proximity to the nucleus (Fig. 4b). We moreover acquired time-lapse data showing the movement of surfRNA in live cells (online supplementary data, see data availability).

**Fig. 4:**
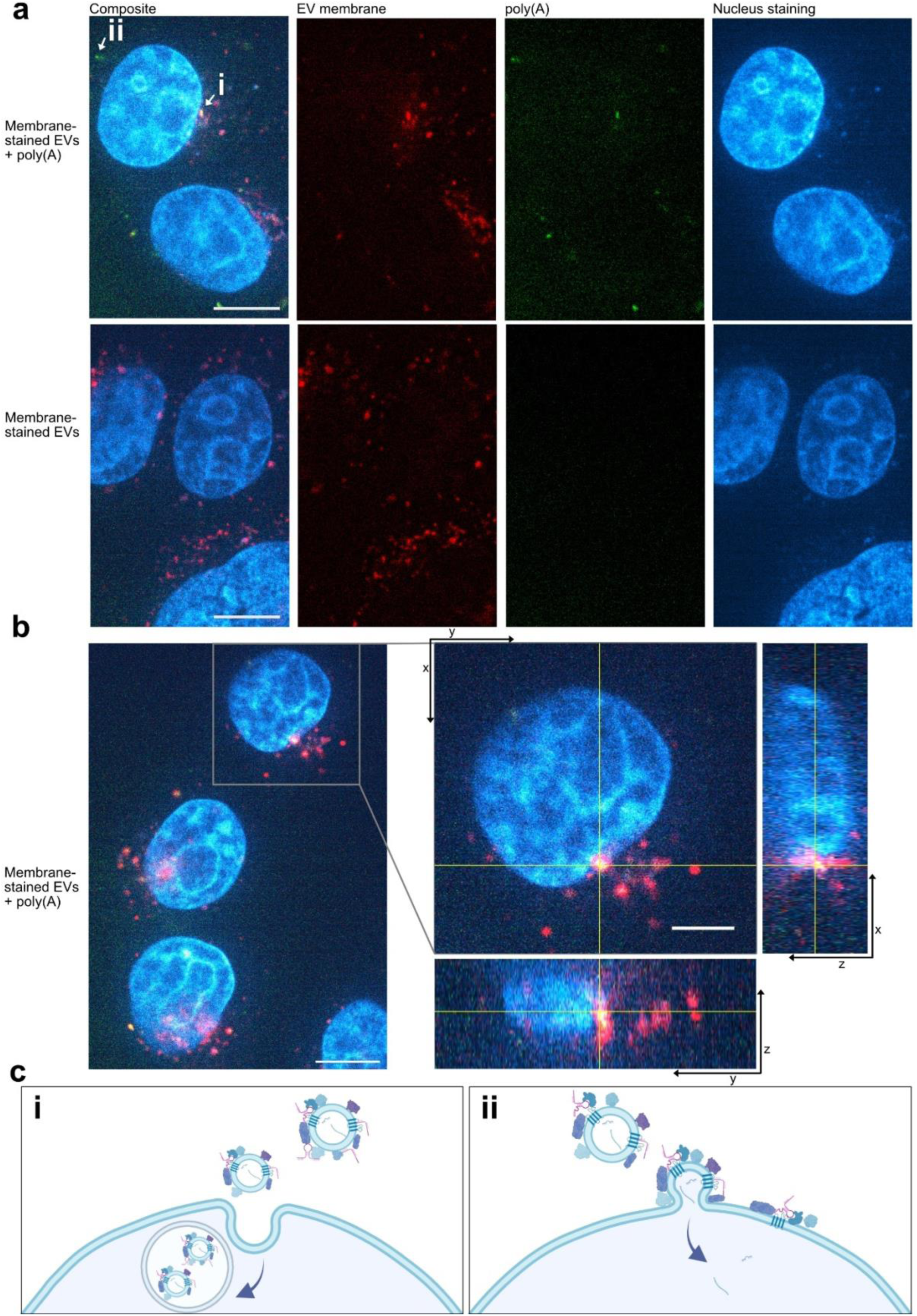
EVs and surfRNAs are co-delivered to target cells and detected in proximity to the nucleus. HeLa cells were incubated with EVs fluorescently labeled for the vesicle membrane (red) and polyadenylated to visualize surfRNA (green). The cell nucleus was stained with DAPI (blue). **a** Representative still images of EVs with and without polyadenylation. Arrows i and ii indicate examples of different EV uptake routes as described in c. Scale bar = 10 µm. **b** Magnification of a cell with surfRNA in the perinuclear region with orthogonal yz and xz images. Magnification scale bar = 5 µm. **c** Schematic of putative EV uptake routes and respective surfRNA fate.

Together, these findings suggest that surfRNAs are co-delivered with EVs through at least two different mechanisms, (i) uptake of surfRNA into cells via endocytosis and (ii) attachment to the cell membrane upon membrane fusion (Fig. 4c).

## Discussion

By providing evidence for the presence of RNA on the surface of EVs, we here introduce the concept of surfRNA. We investigate localization, identity and fate of this novel EV-associated RNA, which is an important step towards understanding the role of RNA in the extracellular and extravesicular space.

Two independent single-vesicle methods were used for direct visualization of EVs and their associated surface RNA. First, we subjected EVs to poly(A) tailing with fluorescent ATP and observed significant labeling compared to Proteinase K and RNase A treated EVs. The utilized enzyme is selective for RNA^30^ and membrane impermeable, which in combination implies presence of RNA on the EV surface. However, the enzyme only labels 3’-ends of RNA and requires a 3’-OH group. It is therefore likely that poly(A) tailing only reveals a subset of the total surfRNA. The application of nanoparticle flow cytometry provided size measurements of particles. Yet, as the polyadenylation causes an increase in size, the sizes detected here are of limited validity.

To detect RNA independent of the accessibility of the strand ends, we then used TOTO-3, a membrane impermeable nucleic acid stain. This indeed resulted in a higher percentage of labeled EVs. Other reasons for the increase in labeled EVs could be the higher sensitivity of TIRF or the presence of surface-bound DNA, as TOTO-3 is not selective for RNA. DNA was previously shown to exist on the EV surface in several studies^44–46^. As even classical EV markers are only found on subsets of vesicles^47–49^, the presence of surfRNA on EVs is likely also limited to a subset of EVs. However, the proportion of EVs with surfRNA could be underestimated in this study, as even TOTO-3 might not stain short, modified or protein-bound RNA.

For TIRF microscopy, the relative particle size, estimated from the membrane signal intensity, showed no correlation with the intensity of the TOTO-3 staining, indicating that larger EVs do not carry more RNA than smaller EVs.

After providing proof for the presence of surfRNA on a single-vesicle level, we investigated the RNA identity by conducting two independent sequencing approaches which aligned in showing that tRFs are the main constituent of surfRNA.

EVs treated with Proteinase K and RNase A could be clearly distinguished from untreated samples by PCA, indicating differences between the RNA inside vs outside the EVs. The relatively smaller distance between treated and lysed samples can be explained by similar ratios of sequences in the samples, although at different amounts of RNA before amplification by PCR.

As described for the poly(A) tailing, the approach where we ligated sequencing adapters to RNA on intact EVs is limited to RNA with an accessible 3’OH group. While this is typically the case for 3’-end of tRNAs and 3’tRFs, the chemistry at the 3’-end of 5’ halves, 5’ fragments and internal fragments depends on the origin and cleavage method of the tRFs^50^. Indeed, we do see 5’ halves and 5’ fragments in both sequencing datasets. 5’ tRFs carrying a 3’ OH group can originate from various cleavage processes, one of them being Dicer-mediated cleavage inside of the cell^51^.

Many of the eleven sequences we found most enriched on the EV surface are linked to disease-relevant cellular functions^52–55^, for instance the response to hypoxia and protection of kidney function^56^. Of note, several of the detected enriched tRFs had already been found in EVs, including a study showing that one of the tRFs could be used as an EV-derived biomarker for Huntington’s disease^57–59^.

Very recently, a sequencing method specifically developed for glycoRNA revealed glycosylated tRNA in EV samples^60^. Although the exact location of the EV glycoRNA was not explored in the study, glycoRNA was originally discovered on the surface of cells, showing its ability to withstand degradation in the extracellular space^21^. It is therefore possible that some of the surfRNA found in our study may be EV glycoRNA. However, as our library preparation excluded RNA with a length of more than 50 nucleotides, our dataset does not contain full length tRNAs. Overall, we present eleven tRNA derived sequences that we detect with high confidence on the EV surface. This could advance our understanding of tRNA biology and related disease and be useful for developing diagnostic tests based on EVs.

Lastly, we directly visualized the interaction of EVs and their surfRNA with target cells by live cell imaging. We observed surfRNA both in proximity to the nucleus and in the periphery of the cell, which is in line with research showing that EVs can either be endocytosed or fuse with the cell membrane^39–41^. The ability of cells to take up free exRNA has recently been underlined^61^. While we do not know yet how and where EVs and surfRNA are attached, surfRNA may bind to EVs in the extracellular space after being released from the producer cell. In that case, some of the events described as cellular uptake of free exRNA could be facilitated by EVs.

While not within the scope of our study, the function of surfRNA will be crucial to understand. An involvement in immune regulation is one possibility, as exRNA can become target of autoantibodies in autoimmune diseases. EVs are known to regulate the immune system, so the presentation of modified RNA on the EV surface could lead to immune tolerance of these exRNAs. However, more research will be necessary to decipher possible functions of surfRNA.

In summary, we introduce and characterize RNA as a new component of the surfaceome of EVs and lay out entry routes into cells. Diagnostics based on tRNA and EVs could profit from a deepened understanding of surfRNA. Further, we hope to inspire research exploring the presence of RNA on synthetic nanoparticles or membrane enveloped viruses, to shed light on RNA in the extracellular space.

## Methods

### Wharton’s Jelly mesenchymal stromal cell culture conditions

WJ-MSC/TERT273 cells (Evercyte GmbH) were passaged twice a week according to the manufacturer’s instructions^25^. In brief, cell culture roux flasks were pre-coated for 2 h at room temperature with fresh Animal component-free cell attachment substrate (ACF, StemCell Technologies, 1:300 dilutions). Mesencult^TM^ ACF Plus medium (StemCell Technologies) was supplemented with Mesencult^TM^ ACF Plus 500x supplement (StemCell Technologies), GlutaMAXTM-I 100X (Gibco) and 200 μg/ml G418 (InvivoGen) and used for cell culturing. Cells were then washed twice with PBS (Gibco) and incubated with CTS^TM^ TrypLE Select Enzyme solution (20 µl/cm2, Gibco) at 37° C until complete detachment from the culture flasks. The cell suspension was then collected in Mesencult medium and centrifuged at 300 x g for 5 min. The cells were cultured in the pre-coated flask in a humidified environment with 5% CO_2_ with a 1:4 to 1:6 split ratio.

### Extracellular vesicle enrichment

For EV enrichment, WJ-MSC/TERT273 cells were cultured in 150 mm cell culture dishes (Corning) for 48 hrs. Upon achieving 70-85% confluency, Mesencult medium was replaced with Opti-MEM reduced serum media (Gibco) and conditioned for 48 hrs. Post 48 hr conditioning, the medium was subjected to differential centrifugation, first at 700 x g for 5 min followed by 2000 x g for 10 min at 4° C to remove cell debri and protein aggregates. To remove particles above 220 nm, pre-cleaned supernatant was filtered through 0.22 μm mPES-filter cups (Merck). The conditioned media was concentrated to 1/10^th^ of the initial volume and diafiltrated using two volumes of initial volume and concentrated to final volume of ∼ 60 ml using TFF system (Repligen) with 300 kDa cut-off hollow fibre filters (MidiKros, 370 cm^2^ surface area, Repligen) at aflow rate of 80 ml/min, a transmembrane pressure of 3.0 psi and a shear rate at 3700 s^-1^, Finally, ∼60 ml of diafiltrate was further concentrated to ∼1 ml using 10 kDa cutt-off Amicon ultra-15 centrifugal filter unit (Merck) at 4°C with 3500 x g. Concentrated EVs were analyzed for size and concentration by nanoparticle tracking analysis (NTA) and 50 µl aliquots were stored in PBS-HAT buffer (^62^ Bobbili et al., 2024) at −80°C for further studies.

### Characterization of EVs

#### Nanoparticle tracking analysis (NTA)

Samples concentrated by TFF were characterized by NTA with a NanoSight NS300 instrument equipped with NTA 2.3 analytical software and an additional 488 nm laser (Malvern panalytical). Samples were diluted to 1:1000 in sterile filtered nuclease free water. Diluted samples were loaded in the sample chamber with camera level 12. Three 60 sec videos were recorded per sample in light scatter mode with 5 sec delays between each recording at 25°C. Screen gain 12, detection threshold 3 were kept constant for all recordings. Using batch process option, all the measurements were analyzed automatically.

#### Western blotting

WJ-MSC cells were collected, and the cell pellet was lysed with 200 µl of RIPA buffer 1x protease/phosphatase inhibitor cocktail (Cell Signaling Technology). Samples were lysed using Bioruptor^®^ (Diagenode), cell lysate was then spun at 12,000 x g for 10 min at 4°C and the supernatant was transferred to a new tube and kept on ice. Protein concentration of the supernatant was quantified by BCA assay (Thermo Fisher Scientific) according to the manufacturer’s instructions. 30 μg of cell lysate and 5E+09 particles were mixed with buffer containing 0.5 M dithiothreitol, 0.4 M sodium carbonate (Na2CO3), 8% SDS, and 10% glycerol, and heated at 95°C for 10 min. The samples were loaded onto NuPAGE Novex 4%-12% Bis-Tris Gel (Invitrogen, Thermo Fisher Scientific) and ran at 120 V in NuPAGE MOPS SDS running buffer (Invitrogen, Thermo Fisher Scientific) for 2 h. The proteins on the gel were transferred to an iBlot nitrocellulose membrane (Invitrogen, Thermo Fisher Scientific) for 7 min using iBlot3 system. The membranes were blocked with 50% Intercept^®^ (TBS) Blocking Buffer (LI-COR) 50% TBS supplemented with 0.1% Tween-20 (Sigma Aldrich) for 1 h at room temperature. After blocking, the membranes were incubated overnight at 4°C with primary antibody solution prepared in 50% Intercept^®^ (TBS) Blocking buffer (TSG101, abcam, ab125011, 1:2000; Syntenin-1, Origene, TA504796, 1:1000; Calnexin, abcam, ab22595, 1:1000). The membranes were washed with TBS supplemented with 0.1% Tween-20 (TBS-T) three times for 5 min and incubated with the corresponding secondary antibody (LI-COR; anti-mouse IgG, 926-68072, 1:10,000; anti-rabbit IgG, 925-32213, 1:10,000) for 1 h at room temperature. Membranes were washed with TBS-T for three time for min, twice with PBS and visualized on the ChemiDoc MP Imaging System (BIO-RAD) at 680 and 800 nm. Images were processed using ImageJ Fuji software.

#### EV surface protein profiling by Multiplex bead-based flow cytometry assay

Surface protein profiling of WJ-MSC EVs was performed via bead-based multiplex EV analysis by flow cytometry (MCASPlex EV IO Kit, human, 130-108-813, Miltenyi Biotec) according to the manufacturer’s instructions. In brief, EV samples containing 2.5E+09 particles/ml were diluted with MACSPlex buffer (MPB) to a final volume of 120 μL and loaded onto wells on a pre-wet and drained MACSPlex 96-well 0.22 μm filter plate. 15 µl of MACSPlex Exosome Capture Beads (containing 39 different antibody-coated bead subsets) were added to each well. The filter plate was then incubated on an orbital shaker overnight (14-16 h) at 450 rpm at room temperature protected from light. To wash the beads, 200 µl of MPB were added to each well and centrifuged 300 x g at room temperature for 3 min. For counterstaining of EVs bound to capture beads with detection antibodies, 135 µl of MPB and 5 µl of each APC-conjugated detection antibody cocktail (anti-CD9, anti-CD63, anti-CD81) were added to each well and the plate was incubated on an orbital shaker at 450 rpm at room temperature for 1 h protected from light. Next, the plate was washed twice by adding 200 µl MPB to each well and centrifuge 300 x g for 3 min at room temperature. This was followed by another wash step with 200 µl of MPB, incubation on an orbital shaker at 450 rpm protected from light for 15 min at room temperature and draining all the wells by centrifugation at 300 x g for 3 min. Subsequently, 200 µl of MPB was added to each well, beads were resuspended by pipetting and transferred to V-bottom 96-well microtiter plate (Thermo Fischer Scientific). Flow cytometry analysis was performed using CytoFLEX S (Beckman Coulter). All samples were automatically mixed and 170 µl of samples were immediately loaded to and acquired by the instrument, resulting in approximately 3,000-5,000 single bead events recorded per well. CytExpert Software (Beckman Coulter) was used to analyze flow cytometric data. Median Fluorescence Intensities (MFI) for all the 39 capture bead subsets were corrected for background by subtracting respective MFI values from matched non-EV buffer (PBS) that were treated as EV-containing samples (buffer + capture beads + antibodies).

#### Transmission electron microscopy (TEM)

WJ-MSC/TERT273–derived extracellular vesicles (EVs) were visualized by transmission electron microscopy, and sample preparation was performed as previously described^63^. Briefly, Formvar-coated copper grids were floated on a 10 µL drop of sample (1 × 10⁹–1 × 10¹⁰ particles, diluted in ddH₂O) for 5 min. Excess liquid was gently removed using Whatman filter paper, after which the grids were rinsed with distilled water and immediately transferred onto a drop of 2% glutaraldehyde for 10 min for fixation. Following an additional brief rinse in distilled water, negative staining was carried out using uranyl acetate replacement stain (UAR_EMS Stain, Electron Microscopy Sciences) by floating the grids on UAR twice for 10 s and once for 60 s, blotting away excess stain after each step. Grids were then air-dried and imaged using a 160 kV FEI Tecnai G2 transmission electron microscope.

### Liposome preparation

Liposomes were prepared from a mixed lipid composition composed of 98 %, 2-Dioleoyl-sn-glycero-3-phosphocholine (DOPC), 0.5 % ATTO-655-1,2-Dioleoyl-sn-glycero-3-phosphoethanolamine (DOPE) (ATTO tec), 0.5 % 1,2-distearoyl-sn-glycero-3-phosphoethanolamine-N-[biotinyl(polyethylene glycol)-2000] (DSPE-PEG(2000) biotin) and 1% 1,2-dioleoyl-sn-glycero-3-phospho-(1’-rac-glycerol) (DOPG) in mole percentage. All lipids were suspended in chloroform and mixed followed by evaporation using N_2_ and vacuumed for 1 hour. The lipid film was hydrolyzed in 200 µl nuclease free water with 400 µM tRNA (Sigma-Aldrich) for liposomes with lumen tRNA and in 200 µl nuclease free water for empty liposomes for 30 min. The liposomes were flash-freezed and thawed for 10 cycles to ensure unilammelarity and extruded at 100 nm.

### Poly(A)-tailing

The surface RNA of EVs was labeled using a poly(A)-tailing kit (Invitrogen). Thirty percent of ATP was replaced by 8-(6-aminohexyl)-amino-ATP-ATTO-647N (Sigma-Aldrich). To avoid unspecific binding of the fluorophore, the overall amount of ATP was reduced to ¼ of the recommended concentration. 5 × 10×8 EVs were used per condition and labeled for 15 min at 37 °C. The EVs were then membrane labeled as described below. For the control with Proteinase K and RNase A treated EVs, the same amount of EVs were incubated with 0.15 U of Proteinase K (NEB) for 45 min at 37 °C. cOmplete^TM^ EDTA-free Proteinase Inhibitor Cocktail (Roche) was added and incubated for 15 min at 37 °C. RNase A (Thermo Fisher) was added at a concentration of 125 µg/ml for 30 min at 37 °C and RiboLock RNAse inhibitor (Thermo Fisher) at 2 U/µl for another 15 min at 37 °C.

### Membrane labeling and biotinylation

The membrane of EVs was labeled using an established method ^64^. EVs were mixed with 300 DOPE-ATTO and per EV in PBS and incubated for 20 min at 37 °C under gentle shaking to allow insertion into the membrane. For TIRF microscopy, EVs were additionally biotinylated in the same step with a concentration of 150 DSPE-biotin molecules per EV. For TIRF microscopy and cell uptake experiments, unbound lipids were washed away using a pre-washed Amicon Ultra centrifugal filter with 50 kDa cutoff (Millipore). Three washing steps with 6200 g were performed and EVs were extracted from the filter with 1200 g.

### TOTO-3 labeling of EVs and liposomes

The dimeric cyanine stain TOTO-3 Iodide is membrane impermeable and stains both RNA and DNA. Membrane labeled and biotinylated EVs were stained with 0.5 µM TOTO-3 for 20 min at room temperature. For the negative control, TOTO-3 was incubated in PBS respectively. Liposomes were treated with RNase A (Thermo Fisher) at a concentration of 125 µg/ml for 30 min at 37 ° before staining with 0.5 µM TOTO-3 for 20 min at room temperature.

EVs and liposomes were washed using a pre-washed Amicon Ultra centrifugal filter with 50 kDa cutoff (Millipore) and measure by TIRF microscopy.

### Nano particle flow cytometry

EVs were analyzed with a CytoFLEX Nano (Beckman Coulter). All samples were filtered through 0.45 µM syringe filters (Cobetter filtration) before measuring and further diluted to yield between 500 and 5000 events per second. Analysis was performed on the samples after removing the background signal based on PBS. For each sample, 100 000 events were recorded. The negative controls with membrane dye and poly(A)-tailing in PBS were diluted equal to the samples and recorded for 1 min at equal flow rates. For the Geometric Mean Intensity (GMI), the background intensity of unstained EVs was subtracted.

### TIRF microscopy

RNA labeling with TOTO-3 was assessed using an Oxford Nanoimager (ONI) microscope. To capture the EVs, a biotinylated lipid-anchor (DSPE-biotin) was inserted in the EV membrane as described above. The biotinylated and fluorescently labeled EVs were captured on passivated glass slides with a microscope chamber (sticky-Slide VI 0.4, ibidi). The glass surface passivation was prepared as described previously ^64, 65^. In brief, they were cleaned, passivated, coated with PLL-PEG and PLL-PEG grafted biotin in a 1:100 ratio and subsequently covered with neutravidin. EVs were added to the surface and allowed to bind for 5 min, before being washed away using 2 ml PBS. The data was acquired with a 100× 1.4 NA oil immersion objective. 162 images were acquired for each condition. Images in two channels were taken successively for each field of view to avoid bleaching, with the 488 laser at 7% and the 640 laser at 15% and an exposure time of 100 ms for EVs. For liposomes, the 488 laser was set to 0.02 % and the 640 laser to 20 % and the exposure time was 100 ms. One field of view corresponds to a physical field of view of 50 μm x 80 μm per channel.

TIRF microscopy data was analyzed using an in-house developed automated python-based image analysis. Spots corresponding to EVs were detected as described before^6465^. The membrane channel was used for spot detection and the background-corrected signal for the respective spots was extracted from both channels (membrane and TOTO-3). For each experiment, the threshold for positive vs negative TOTO-3 signal was set based on the standard deviation of the TOTO-3 signal in the unstained condition multiplied by 2.5. For better comparability, the threshold for liposomes was set to 2.27 as in the presented EV dataset in Fig. 2c. An equal number of particles was used for the three conditions within each experiment, based on the condition with lowest spot count.

### Small RNA sequencing

#### Comparison of untreated and surface-treated EVs

EVs enrichment, immunoprecipitation and RNA isolation was described previously ^37^. Briefly, immunoprecipitation was performed using a hCD81 exosome isolation kit (Miltenyi biotec) or µMACS^TM^ HA Isolation Kit (Miltenyi biotec). EV samples were treated with Proteinase K and RNase A as described for poly(A) tailing. For EV lysis and intraluminal Proteinase K and RNase treatment, EVs were incubated in 0.3 % Triton X for 15 min at room temperature, followed by Proteinase K and RNase A treatment as described. Untreated samples were kept at same temperature conditions without treatment.

RNA was isolated using the miRNeasy Mini Kit (Qiagen) and RNeasy Min Elute Cleanup Kit (Qiagen) and RNA integrity was confirmed on an Agilent 2100 Bioanalyzer. RNA libraries were constructed using the RNA Library Prep kit (Qiagen) by adapter ligation on both ends of the RNA, reverse transcription, addition of unique molecular identifiers (UMI) and amplification. Library quality was assessed on the Agilent2100 Bioanalyzer and by qPCR using KAPA Library Quantification kit (Roche). Equal amounts of libraries were pooled and sequencing was performed on an Ilumina Novaseq.

#### Comparison of surface-ligated EV RNA and isolated EV RNA

TFF-isolated EVs were used for specific sequencing of surfRNA. Total EV RNA was isolated using the miRNeasy Serum/Plasma Advanced Kit (Qiagen, Cat. No. 217204) following the manufacturer’s protocol. RNA was eluted in 12 µL of RNase-free water and quantified using the Bioanalyzer RNA 6000 Pico Kit (Agilent, Cat. No. 5067-1513).

For small RNA library preparation, 5 µL of eluted RNA was processed using the QIAseq miRNA Library Kit (Qiagen, Cat. No. 331505). Briefly, 3’ and 5’ adapters were ligated to the RNA, followed by reverse transcription with a primer containing a unique molecular index (UMI). Library amplification was performed using 24 PCR cycles to introduce the sample index.

For EV surface miRNA library preparation, EVs were directly subjected to 3’ adapter ligation by incubation at 16 °C for 16 h. RNA was then extracted, and library construction was subsequently continued according to the standard protocol.

Library quality and quantity were assessed using the Bioanalyzer High Sensitivity DNA Analysis Kit (Agilent, Cat. No. 5067-4626). Finally, libraries were pooled and sequenced on one lane of a NovaSeq Plus 10B flow cell (Illumina, San Diego, CA, USA) using single-end 150 bp sequencing.

#### Sequencing data analysis

For both datasets, mapped reads were assigned to annotated RNAs depending on the RNA class and further analyzed using R (version 2025.05.1). For the dataset based on surface-treated vs untreated EVs, differential expression analysis was performed using DESeq2. Significance was assigned when Benjamini-Hochberg adjusted p-value < 0.05 and log_2_ fold change ≥ 1.

In the dataset based on EV external adapter ligation, counts of zero were replaced with 1 to allow ratio calculations. The ratio of counts in the surface-aligned sample to the whole EV sample was then calculated for each tRF and the top 100 fragments were retained for comparison.

Overlap between the datasets was assessed after trimming all sequences to the tRF type and codon, and identically for the untrimmed sequences.

### Uptake experiments and spinning disc confocal imaging

For uptake experiments, HeLa cells were cultured in ibidiTreat µ-Slides VI 0.4 (ibidi). 12 000 cells were seeded one day prior to imaging. EVs were labeled by poly(A) tailing as described and 4 x 10^9^ EVs were added per channel after medium exchange to live cell imaging solution (Invitrogen). The cells were incubated for 30 min at 37 °C before acquisition. Cell nuclei were stained with DAPI as recommended by the supplier. Imaging was performed using an IX83 inverted spinning disk confocal microscope (Olympus) with a 60 x magnification water objective. Laser settings were 3 % for the 488 nm laser and 7 % for the 640 nm laser, the exposure time was 100 ms. The microscope was equipped with an incubation chamber, and all live-cell experiments were performed at 37 °C and 5% CO_2_. Prior to the experiments the microscope was pre-heated.

### Statistical analysis

Statistical significance was assessed using GraphPad Prism 10 or python, respective tests are indicated in the figure description. Significance levels were * = <0.05, ** = <0.01, ***= <0.001, **** = <0.0001.

### Data availability

The authors declare that the data supporting the findings of this study are available within the paper and its Supplementary information files. Additionally, underlying data, source data, sequencing data and code will be made available upon acceptance. Live imaging recordings of EVs in cells can be accessed under https://anon.erda.au.dk/sharelink/eSPKJR9XMp

### Contributions

H.W. performed nanoparticle flow cytometry, TIRF microscopy and live cell imaging. M.B. isolated and characterized EVs. M.G.M produced liposomes and was involved in designing TIRF microscopy and live cell imaging experiments. E.A. and M.B. performed transmission electron microscopy. H.W., Y.Y., M.B. and M.G. were involved in planning, conducting and analyzing the sequencing experiments. H.W., M.G.M. J.G. and J.K. designed the project. J.K. acquired funding for the project. H.W. wrote the manuscript with input from all authors.

## Acknowledgments

This work was supported by the Lundbeck Foundation grant No. R380-2021-1393 for M.G.M, and the Novo Nordisk Foundation grant NNF23OC0081177 (RNA-META) for H.W. We would like to acknowledge Omiics for a good collaboration on the sequencing experiments. In addition, we would like to acknowledge BOKU core facilities for Biomolecular and Cellular Analysis (BMCA) as well as the Bioimaging Core Facility, Aarhus University and the FACS Core Facility, Aarhus University for use of equipment and support and the Carlsberg Foundation, grant CF23-1020 for providing the CytoFLEX Nano (Beckman Coulter).

## Notes

https://anon.erda.au.dk/sharelink/eSPKJR9XMp

### Competing Interest Statement

The authors have declared no competing interest.

