## Supplementary Information for "Native extracellular vesicles display surface bound RNAs that are co-delivered to cells"

SUPPLEMENETARY INFORMATION:

Table of contents:

Supplementary Figure S1: Characterization of EVs

Supplementary Figure S2: Polyadenylation of surfRNA

Supplementary Figure S3: TOTO-staining of EVs

Supplementary Figure S4: TOTO-staining of liposomes

Supplementary Figure S5: Small RNA sequencing

Supplementary Figure S6: Live cell imaging of surfRNA delivery to cells

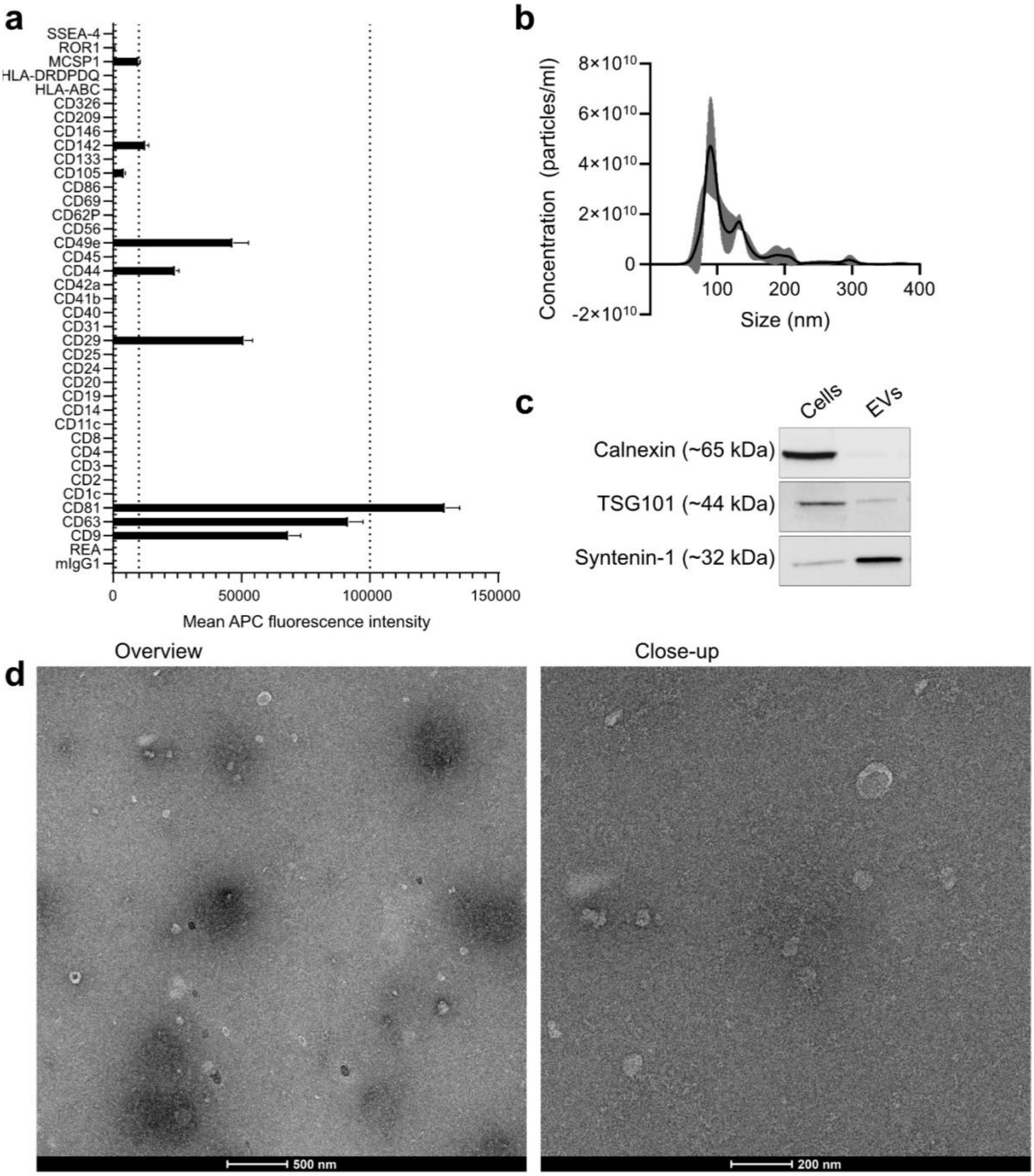


**Supplementary Fig. S1: Characterization of TFF-enriched EVs. a** Multiplex bead-based flow cytometry showing the expression of EV surface protein profile. **b** Nanoparticle tracking analysis showing the size and concentration of EVs. **c** Western blots on eluates of EVs and producing cells for the EV marker proteins syntenin-1 and TSG101 and non-EV marker calnexin (ER specific). **d** Representative TEM images of EVs. Scale bar overview: 500 nm. Close up: 200 nm.

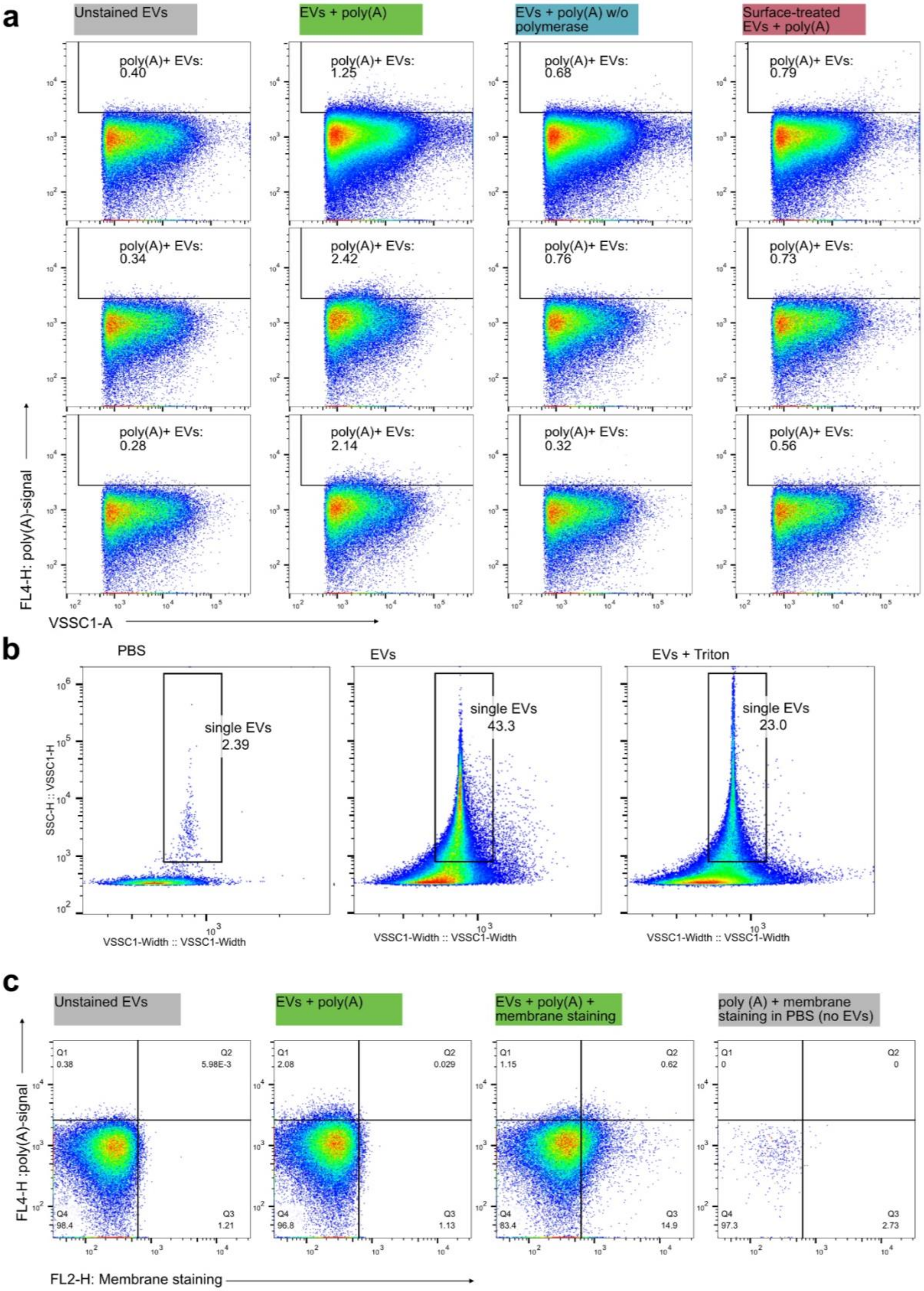


**Supplementary Fig. S2: Polyadenylation of surfRNA. a** Scatterplots of EVs with polyadenylated surface RNA and experimental controls for three independent experiments. The bottom row is presented in Fig. 1b. **b** Gating strategy and controls. All data shown in Fig.

1 only include data points from the single EV populations. Triton lysis shows a reduction of EVs and a shift towards smaller particles. **c** Representative scatterplots (n=2) showing poly(A) tailing of EVs in combination with membrane staining.

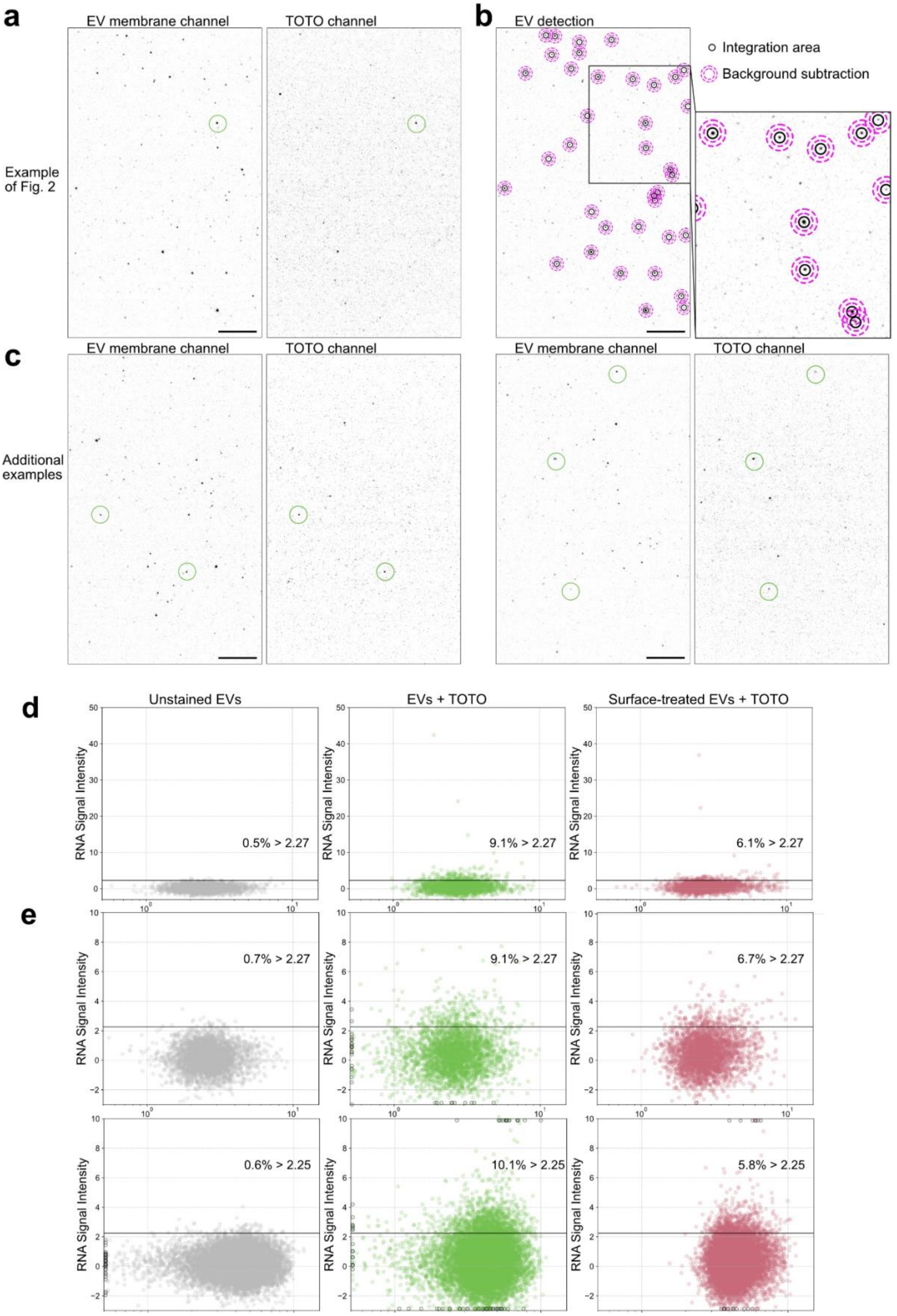
a
EV membrane channel
TOTO channel
Example of Fig. 2
b
EV detection
Integration area
Background subtraction
c
EV membrane channel
TOTO channel
Additional examples
EV membrane channel
TOTO channel
d
Unstained EVs
EVs + TOTO
Surface-treated EVs + TOTO
RNA Signal Intensity
0.5% > 2.27
9.1% > 2.27
6.1% > 2.27
e
0.7% > 2.27
9.1% > 2.27
6.7% > 2.27
0.6% > 2.25
10.1% > 2.25
5.8% > 2.25

**Supplemenetary Fig. S3: TOTO-staining of EVs. a** Full field of view of images shown in Fig. 2e. **b** Field of view shown in c with visualized automated EV detection. The intensity of the area within the black circle is integrated and the median intensity of the area between the pink circles is subtracted for local background correction. **c** Additional representative field of views showing the EV membrane and TOTO-3 signal. Scale bar = 10 µm. **d** Extended scatterplot shown in Fig. 2e showing the percentage of TOTO-stained EVs. **e** Scatterplots showing the percentage of TOTO-stained EVs of experiments n2 and n3. Thresholds are 2.5 x standard deviation of red intensity in the unstained condition. Number of analyzed EVs: 3360 and 12260. Datapoints that were outside the axis range are depicted with black outline on the respective graph edge.

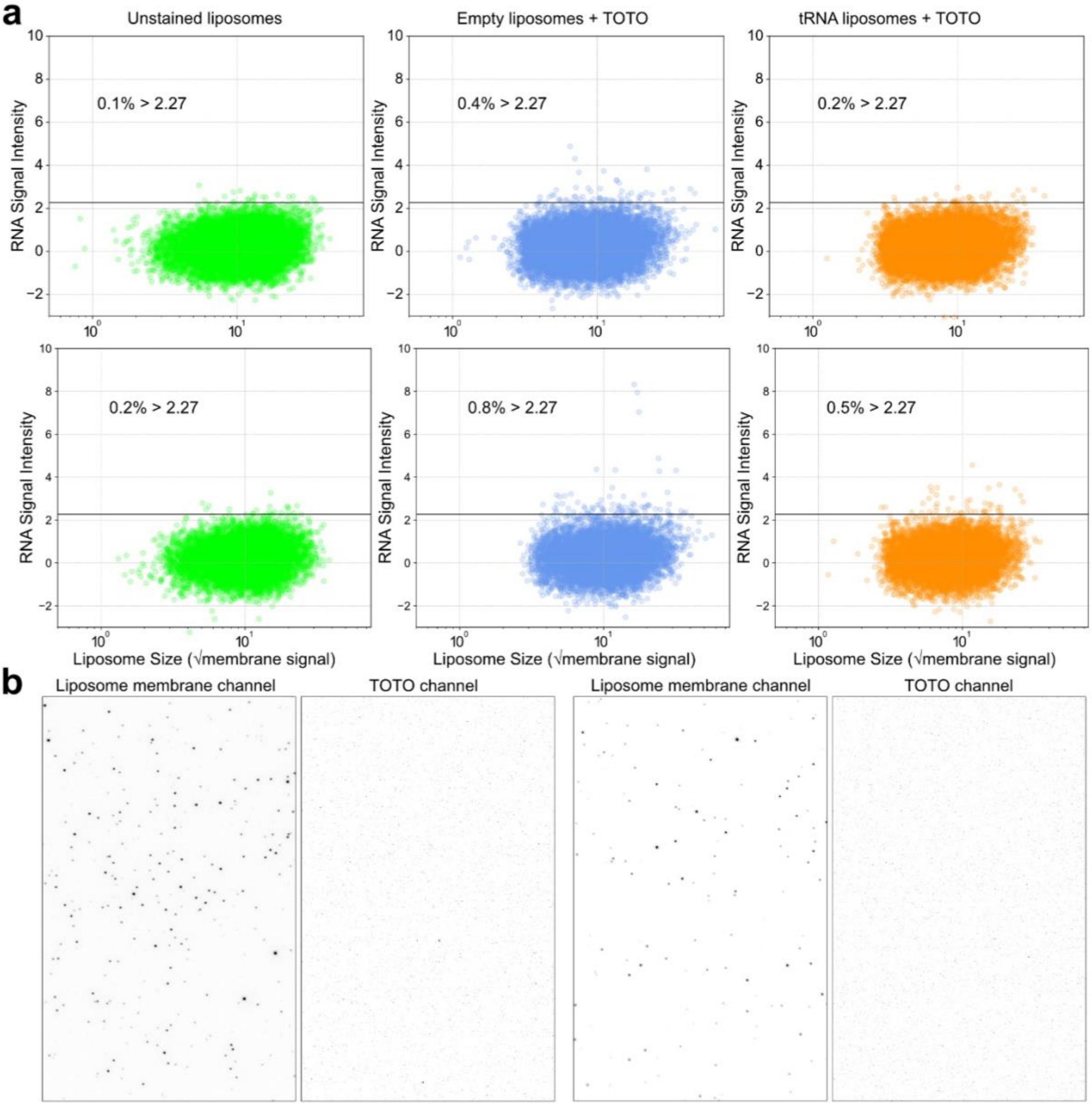


**Supplementary Fig. S4: TOTO-3 staining of liposomes. a** Scatterplots showing the percentage of TOTO-stained liposomes in two experiments. Identical threshold to the scatterplot presented in Fig. 2e. **b** Representative field of views showing the liposome membrane signal and TOTO-3 signal. Scale bar = 10 µm.

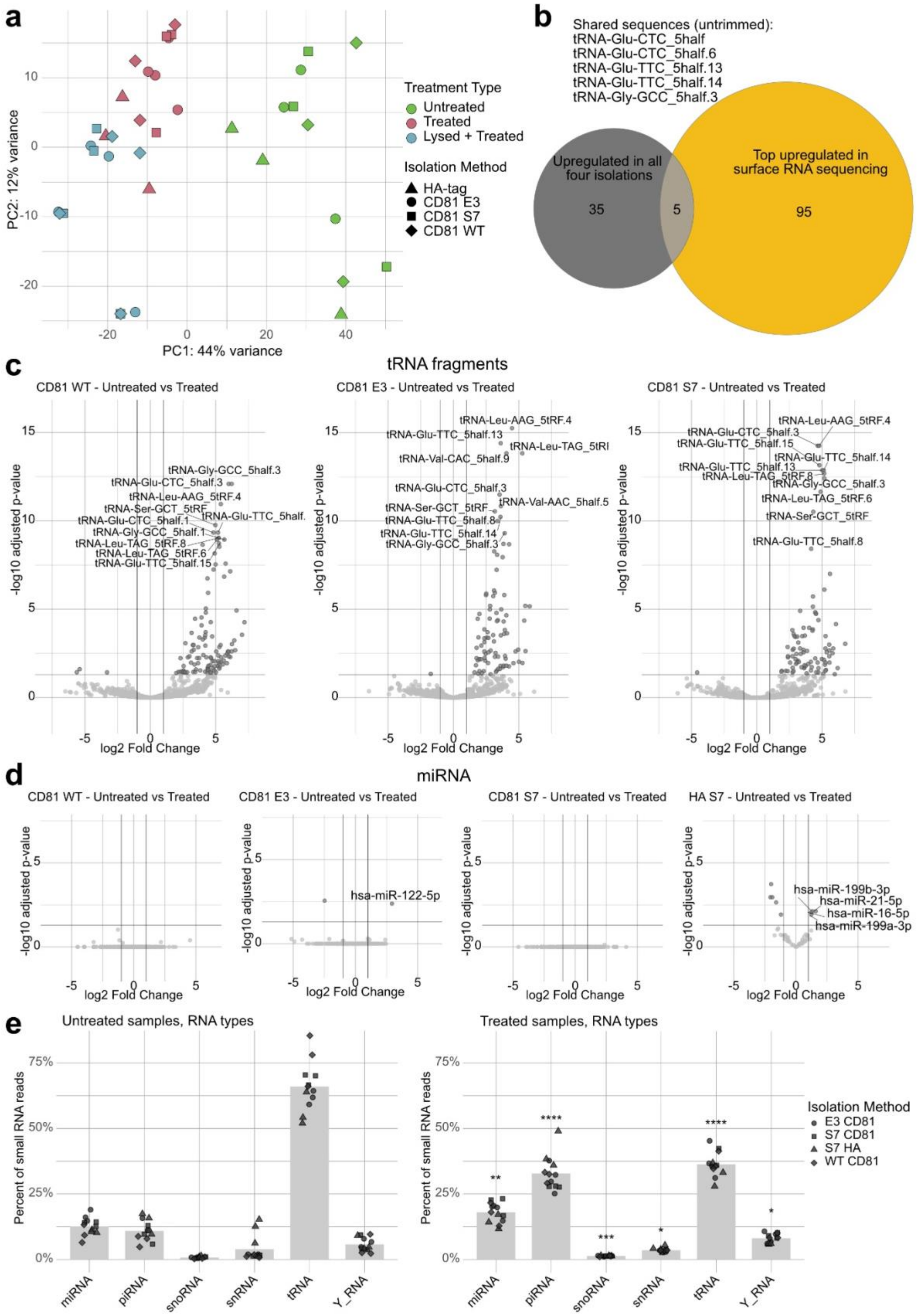


**Supplementary Fig. S5: Small RNA sequencing. a** Principal component analysis (PCA) of variance-stabilized tRNA fragments of untreated EVs, surface-treated EVs (Proteinase K + RNase A) and lysed and treated EVs. Different shapes indicate isolation methods. **b** Euler diagram of untrimmed sequences, overlap of tRNA fragments that were upregulated in all four isolations of Fig. 3c with the 100 most upregulated sequences of the selective surface RNA sequencing illustrated in Fig. 3d. **c** Vulcano plots for tRNA fragments of EVs isolated by indicated methods, each comparing untreated and surface-treated EVs. Significance was defined as FDR < 0.05 and $|\log_2$ fold change$| > 1$. The 10 sequences with the lowest adjusted p-value are labeled. **d** Vulcano plots for miRNAs of EVs isolated by indicated methods, each comparing untreated and surface-treated EVs. Significance was defined as FDR < 0.05 and $|\log_2$ fold change$| > 1$. The 10 sequences with the lowest adjusted p-value are labeled if significance was reached. **e** Changes in relative amount of RNA types after Proteinase K and RNase A treatment. Statistical significance: Wilcoxon test comparing percentages of RNA in untreated and treated samples.

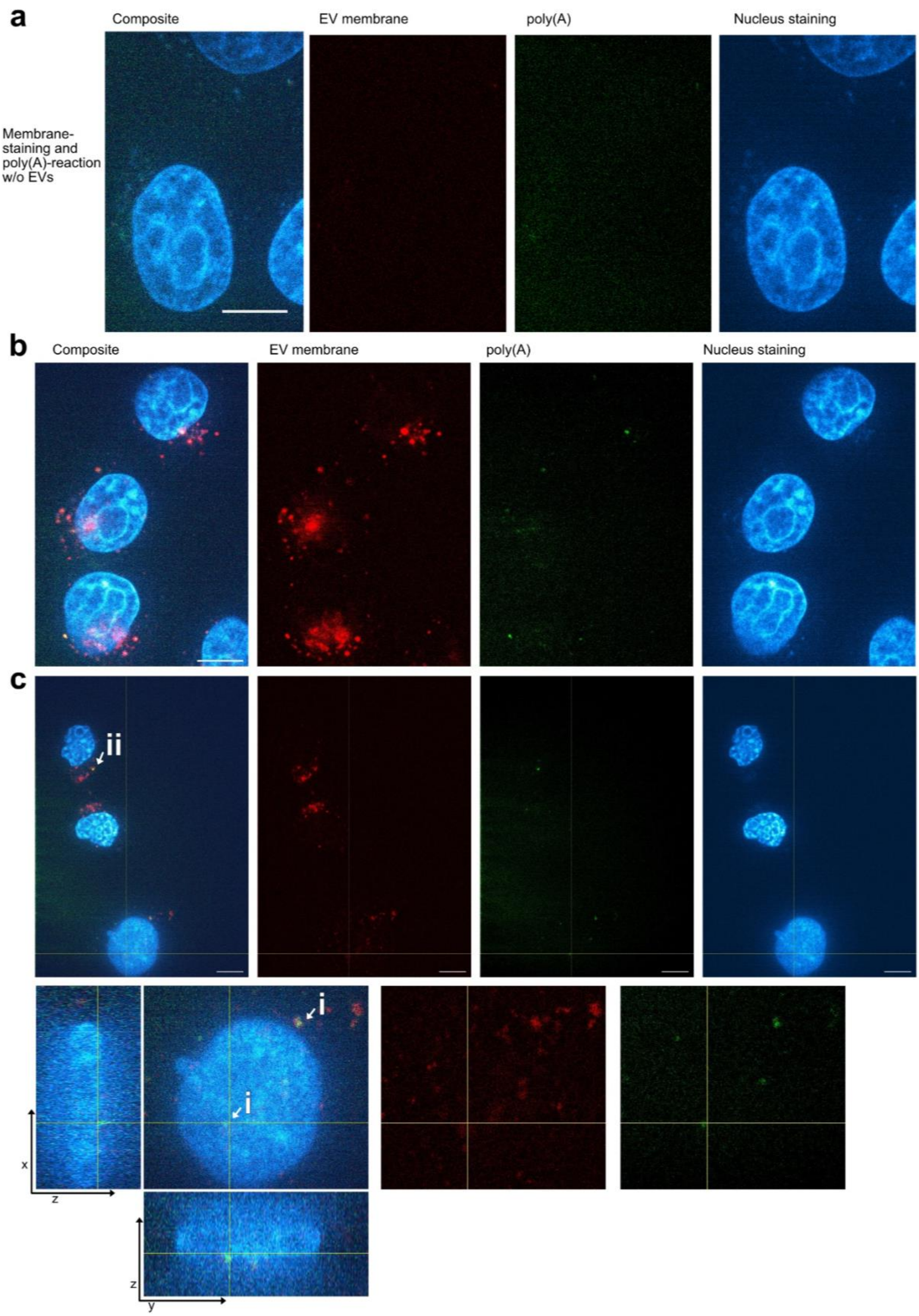


**Supplementary Fig. S6: Live cell imaging of surfRNA delivery to cells.** Vesicle membrane (red) and polyadenylated surfRNA (green). The cell nucleus was stained with DAPI (blue). **a** Negative control consisting of membrane dye and poly(A)-tailing reaction performed in PBS without EVs. Scale bar = 10 μm. **b** Images of separate channels for Fig. 4b. **c** Additional field of view with close-up showing surfRNA in nuclear proximity and in the cellular periphery.
